# Carbon starvation-induced lipoprotein Slp directs the synthesis of catalase and expression of OxyR regulator to protect against hydrogen peroxide stress in Escherichia coli

**DOI:** 10.1101/386003

**Authors:** Xiaoxia Li, Yuanhong Xie, Junhua Jin, Hui Liu, Xiuzhi Gao, Lixia Xiong, Hongxing Zhang

## Abstract

*Escherichia coli* can induce a group of stress-response proteins, including carbon starvation-induced lipoprotein (Slp), which is an outer membrane lipoprotein expressed in response to stressful environments. In this paper, *slp* null mutant

*E. coli* were constructed by insertion of the group II intron, and then the growth sensitivity of the *slp* mutant strain was measured under 0.6% (vol/vol) hydrogen peroxide. The changes in resistance to hydrogen peroxide stress were investigated by detecting antioxidant activity and gene expression in the *slp* mutant strain. The results showed that deletion of the *slp* gene increased the sensitivity of *E. coli* under 0.6% (vol/vol) hydrogen peroxide oxidative stress. Analysis of the unique mapping rates from the transcriptome libraries revealed that four of thirteen remarkably up/down-regulated genes in *E. coli* were involved in antioxidant enzymes after mutation of the *slp* gene. Mutation of the *slp* gene caused a significant increase in catalase activity, which contributed to an increase in glutathione peroxidase activity. The *katG* gene was activated by the OxyR regulator, which was activated directly by 0.6% (vol/vol) hydrogen peroxide, and HPI encoded by *katG* was induced against oxidative stress. Therefore, the carbon starvation-induced lipoprotein Slp regulates the expression of antioxidant enzymes and the transcriptional activator OxyR in response to the hydrogen peroxide environment, ensuring that cells are protected from hydrogen peroxide oxidative stress at the level of enzyme activity and gene expression.

## Introduction

In nature, bacteria are often subjected to adverse environmental conditions, such as high temperature, osmotic stress or oxidative stress [1, 2]. When non-differentiating bacteria such as *E.coli* are exposed to environmental stress, it could induce the synthesis of more than 50 different types of stress-response proteins, and the accumulation and degradation rates of induced partial proteins have a great effect on cell survival, including the Slp protein [3]. The *slp* gene is located at 78.6 centrosomes on the *Escherichia coli* genetic map, and the −10 and −35 regions upstream of the mRNA start site are characteristic of the σ^70^ promoter. Comparison of the *slp* region to *E.coli* sequences in the GenBank database indicated that this carbon starvation-inducible gene encoded a protein with an unprocessed molecular mass of 25 kDa [4]. Slp protein was localized in the outer membrane fraction since it remained in the pelleted membrane fraction following solubilization of the cytoplasmic membrane proteins with 1% Triton X-100 [5]. Bacterial lipoproteins contain an *N*-terminal cysteine residue, which is linked to diglyceride through the thiol group and to a fatty acid on the terminal amino group, consistent with the Slp protein having such an *N*-terminal modification, and the protein was resistant to standard Edman degradation; these criteria indicate that the *slp* gene encodes a new lipoprotein [4, 6]. Early studies found that *E. coli* regulation of the reverse environment mainly depends on the following three notions: the cAMP-dependent protein involved in the absorption and utilization of alternative carbon sources to ensure cell survival, termed as Cst [7]; and the induced expression mechanism of Pex protein was similar to the generalized resistance state of the cell survival characteristics in the stationary phase [1, 8]. Expression of Slp protein during carbon starvation and the stationary phase was dependent on neither cAMP/CRP nor σ^S^ [9].

In recent years, it has been found that the regulation of oxidative stress in *E. coli* is mainly mediated by alkyl peroxide reductase AhpCF and the OxyR stress regulator as well as the catalase family [10]. Alkyl peroxide reductase is a highly efficient two-component disulfide oxidoreductase that relies solely on the AhpC subunit (22 kDa) and the AhpF subunit (52 kDa) for the reduction of bacterial intracellular peroxide substrates [11]. The AhpF subunit transfers electrons via NADH and specifically catalyzes the non-peroxidase AhpC subunit to convert low concentrations (less than 20 μM) of hydrogen peroxide and organic peroxides into water, to protect cells against oxidative stress [12]. However, OxyR, a positive regulator of hydrogen peroxide-inducible protein, can be activated by increases in the hydrogen peroxide concentration or decreases in the thiol/disulfide ratio, and the OxyR regulator senses low concentrations of peroxidation and induces the decomposition of hydrogen peroxide when *E. coli* is grown in an oxidative stress environment of hydrogen peroxide [13, 14]. At the same time, the catalase family HPI (*katG*) and HPII (*katE*) also plays an indispensable role in oxidative stress [10, 15]. Bifunctional catalase/peroxidase (HPI) was encoded by *katG* and monofunctional catalase (HPII) encoded by *katE*, all of which are heme-containing enzymes involved in the dismutation of H_2_O_2_ into O_2_ and H_2_O [15]. In response to low molar concentrations of H_2_O_2_, OxyR regulator protein, which controls the expression of the *katG* gene, is activated during environmental stimulation. Further HPI (*katG*) was induced by transcription during logarithmic growth [16]. HPII is not peroxide inducible, and its gene, *katE*, is transcribed by RNA polymerase containing the alternative σ^S^ [10].

In this paper, the *slp* gene of mutant *E. coli* was constructed by the Targetron technique, and the sensitivity of the *slp* mutant strain to hydrogen stress was investigated to explore the regulation mechanism of Slp on oxidative stress in *E. coli*.

## Materials and methods

### Strains, media, and cultivation conditions

*E. coli* BL21(DE3) wild-type (F^−^ompT hsdS_B_(r_B_^−^ m_B_^−^) gal dcm(DE3) pLysS Cam^r^) was grown in Luria-Bertani (LB) medium at 37°C with aeration at 200 rpm in ambient atmosphere. When necessary, kanamycin (Kan) (Sigma) and chloramphenicol (Chl) (Sigma) were added at concentrations of 50 μg/ml and 25 μg/ml, respectively. Recombinant *E. coli* strains were selected and maintained on 25 μg/ml chloramphenicol where indicated.

### Construction of *slp* null mutant *E. coli*

The Targetron plasmid pADC4k-C was used to disrupt the genes in *E. coli* BL21(DE3) according to the Targetron™ Gene Knockout System Kit protocol (Sigma). Group II intron sequences of the gene encoding the carbon starvation-induced lipoprotein Slp (CAQ33824.1) were amplified using the primers listed in Table 1 as designed by sigma-aldrich.com/targetronaccess. Amplification of the Group II intron was performed with a PCR thermocycler (Bio-Rad Co., America) with the following program: 1 cycle at 95°C for 30 seconds, followed by 34 cycles at 95°C for 30 seconds, 55°C for 30 seconds and 72°C for 45 seconds, ending with one final elongation step at 72°C for 5 minutes. The PCR products were cleaved with *Hin*dIII (20 U/ml) and *Bsr*GI (10 U/ml) restriction endonucleases and ligated to the *Hin*dIII and *Bsr*GI sites of the pADC4k-C linearized plasmid to construct the plasmid pADC4k-C-slp. The pADC4k-C-slp plasmid was thermally transferred into wild-type *E. coli* BL21(DE3) using a heat shock method [17]. The transformation reaction was carried out at 37 °C in LB containing 25 mg/ml chloramphenicol and 1% glucose with shaking overnight. Forty microliters of overnight culture was added to 2 ml of LB broth containing 25 µg/ml chloramphenicol and 1% glucose and grown to an optical density at 560 nm of 0.2 at 37 °C, and then the incubator was cooled to 30 °C. Bacterial solution containing 0.5 mM IPTG was incubated at 30 °C for 30 minutes with shaking and grown overnight at 30 °C. The *slp* null mutant *E. coli* was selected and named *E. coli* BL21(DE3)^∆slp^.

**Table 1.**
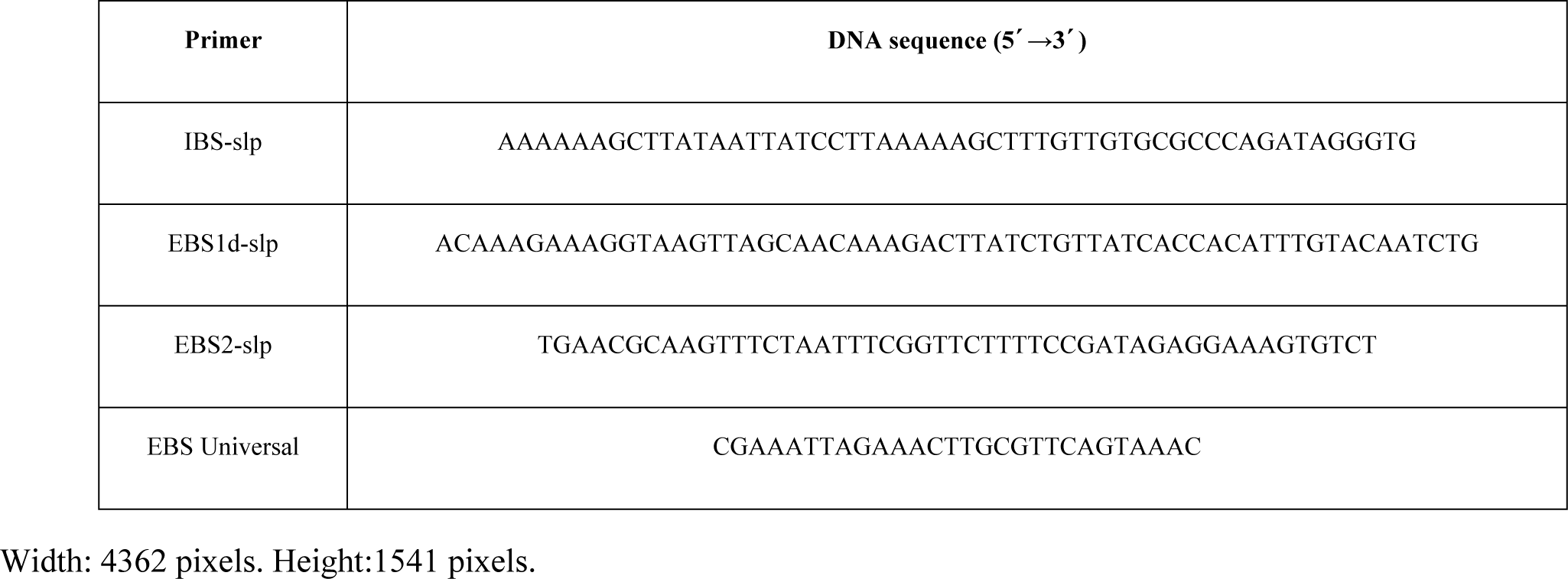
Names and sequences of the *slp* gene mutation primers

### Identification of *slp* null mutant strains

The *E. coli* BL21(DE3)^∆slp^ and wild-type *E. coli* BL21(DE3) were inoculated into LB medium with an initial concentration of 10^9^ CFU/ml and cultured at 37 °C for 12 hours. The genomic DNA of *E. coli* was isolated using a TIANamp Bacteria DNA Kit (Tiangen Biotech Co., Beijing, China) according to the manufacturer’s instructions. The mutations of the *slp* gene insertion site were verified by PCR using the primers slpF (5‘-TGGGGTGGAAATAGAAATAC-3’) and slpR 5‘-GCCGCTTCAGGCACAGAG-3’) at the intron-gene linker with the following program: 1 cycle at 95°C for 30 seconds, followed by 34 cycles at 95°C for 30 seconds, 58°C for 30 seconds and 72°C for 1 minute, ending with one final elongation step at 72°C for 5 minutes.

The expression of *slp* in *E. coli* was also examined. The total RNA of *E. coli* was extracted using a TIANamp total RNA extraction kit (Tiangen Biotech Co., Beijing, China) according to the manufacturer’s instructions. The cDNA was synthesized using a PrimeScript™ 1st Strand cDNA Synthesis Kit (Takara, Dalian, China). Real-Time PCR was used to analyze the transcript expression of the *slp* gene in *E. coli* using the primers yzF (5´-CACTCATCCTCAGCCTTTCAT-3´) and yzR (5´-CGGTAATACAGCGATTTCTAACA-3´). Amplification of the *slp* gene was performed using the 7500 Fast Real-Time PCR System (Applied Biosystems) with the following program: 1 cycle at 95 °C for 20 seconds, and 40 cycles at 95 °C for 15 seconds, 60 °C for 1 minute and 72 °C for 30 seconds. The amplification reactions were carried out with a real-time PCR Premix for SYBR Green II (takara) in a final volume of 20 μl. Quantification of the tested gene expression was done using the comparative 2(-Delta Delta C(T)) method [18].

### Generation of Growth Curves under hydrogen peroxide stress

Subcultures of *E. coli* were grown overnight in LB medium at 37 °C. Then, the *E. coli* BL21(DE3)^∆slp^ and wild-type *E. coli* BL21(DE3) were inoculated with an initial concentration of 10^9^ CFU/ml in LB medium containing 0.6% (vol/vol) H_2_O_2_ and cultured at 37°C for 3 hours. The OD_600_ (optical densities 600nm) of cultures were measured every half hour, and the mean value of the triplicate cultures was plotted. All experiments were repeated three times.

### The RNA-sequencing of transcriptome libraries

*E. coli* BL21(DE3)^∆slp^ and *E. coli* BL21(DE3) were grown in LB medium containing non-hydrogen peroxide or 0.6% (vol/vol) H_2_O_2_ with an initial concentration of 10^9^ CFU/ml and cultured at 37 °C for 3 hours. Total RNA from 3 biological replicate samples (Wild-type *E. coli* BL21(DE3) of the C1 group grown in a non-hydrogen peroxide environment; *E. coli* BL21(DE3)^∆slp^ of the C2 group grown in a non-hydrogen peroxide environment; Wild-type *E. coli* BL21(DE3) of the C3 group grown in a 0.6% (vol/vol) H_2_O_2_ environment; *E. coli* BL21(DE3)^∆slp^ of the C4 group grown in a 0.6% (vol/vol) H_2_O_2_ environment was extracted from each group, and the ribosomal RNA from the total RNA was removed by the Ribo-Zero rRNA Removal Kit (Epicentre Biotech Co., Germany). The total RNA Library was constructed using a TruSeqTM Stranded Total RNA Library Prep Kit for the experiment. After purification of the cDNA using the MinElute PCR Purification kit (Qiagen Biotech Co., Germany), bands of approximately 150-200 bp were recovered. The cDNA was amplified with 15 PCR cycles and quantified withTBS380 (Picogreen). Clusters were generated by bridge PCR on the cBot. A cDNA library was sequenced on an Illumina HiSeqTM 2000 using Illumina’s protocol (IGENCODE Technology Co., Ltd (Beijing)). The details of the methods for alignment of the read and quantitation of gene expression were represented according to the method [19].

Quality control of the microarray was performed using Bead Studio (Illumina Genome Studio Software), and microarray data normalization on the median and background was conducted. Microarray statistical analyses were performed by Biometric Research Branch array tools (http://linus.nci.nih.gov/BRB-ArrayTools.html) and the Multi Experiment Viewer software [20]. General filters were applied before analysis, and probes were excluded if the percentage missing exceeded 50%. If they had a fold difference >2.0, if the univariate p-value was<0.05, and if the false discovery rate (FDR) was<0.001, then genes were identified as differentially expressed.

### Antioxidase activity assays

*E. coli* BL21(DE3)^∆slp^ and wild-type *E. coli* BL21(DE3) were grown in LB medium containing 0.6% (vol/vol) H_2_O_2_ with an initial concentration of 10^9^ CFU/ml and cultured at 37 °C for 3 hours. Ten milliliters of the fermentation were taken at intervals of 0.5 hours. After centrifugation at 10,000 rpm for 5 minutes, the pellets were re-suspended in 20 mM Tris–HCl buffer, pH 8.5, and lysed by a Mini-BeadBeater-16 (BioSpec products) grinding beads homogenizer for 2×15 seconds. The lysate was then centrifuged at 10,000 rpm for 5 minutes, the supernatant was filtered through a 0.22μm filter and was stored at −80 °C [21]. With bovine serum albumin (BSA) as a standard protein, the protein content of the samples was determined by the Micro BCA method Protein Quantitative Detection Kit C503061 (Sangon Biotech (Shanghai) Co., Ltd).

Antioxidase activities of *E. coli* were examined using the relevant kits (Nanjing Bioengineering Research Institute, Nanjing, China). Mainly, the catalase activities of the samples were determined with the visible light method using the catalase detection kit (A007-1). The glutathione S-transferase and peroxidase activity of samples were determined with the visible light method using the glutathione S-transferase detection kit (A004) and glutathione peroxidase detection kit (A005). The superoxide dismutase activity of the samples was determined with the hydroxylamine method by using the superoxide dismutase detection kit (A001-1). The malondialdehyde content of the samples was examined with the thiobarbituric acid method by using the malondialdehyde detection kit (A003-2). All of these experiments were repeated three times.

### Detection of oxidative stress gene expression

The *E. coli* BL21(DE3)^∆slp^ and wild-type *E. coli* BL21(DE3) were grown in LB medium containing 0.6% (vol/vol) H_2_O_2_ with an initial concentration of 10^9^ CFU/ml and cultured at 37 °C for 3 hours. The extraction of total RNA and synthesis of cDNA were performed as previously described. Primers used for real-time PCR are listed in Table 2.

**Table 2.**
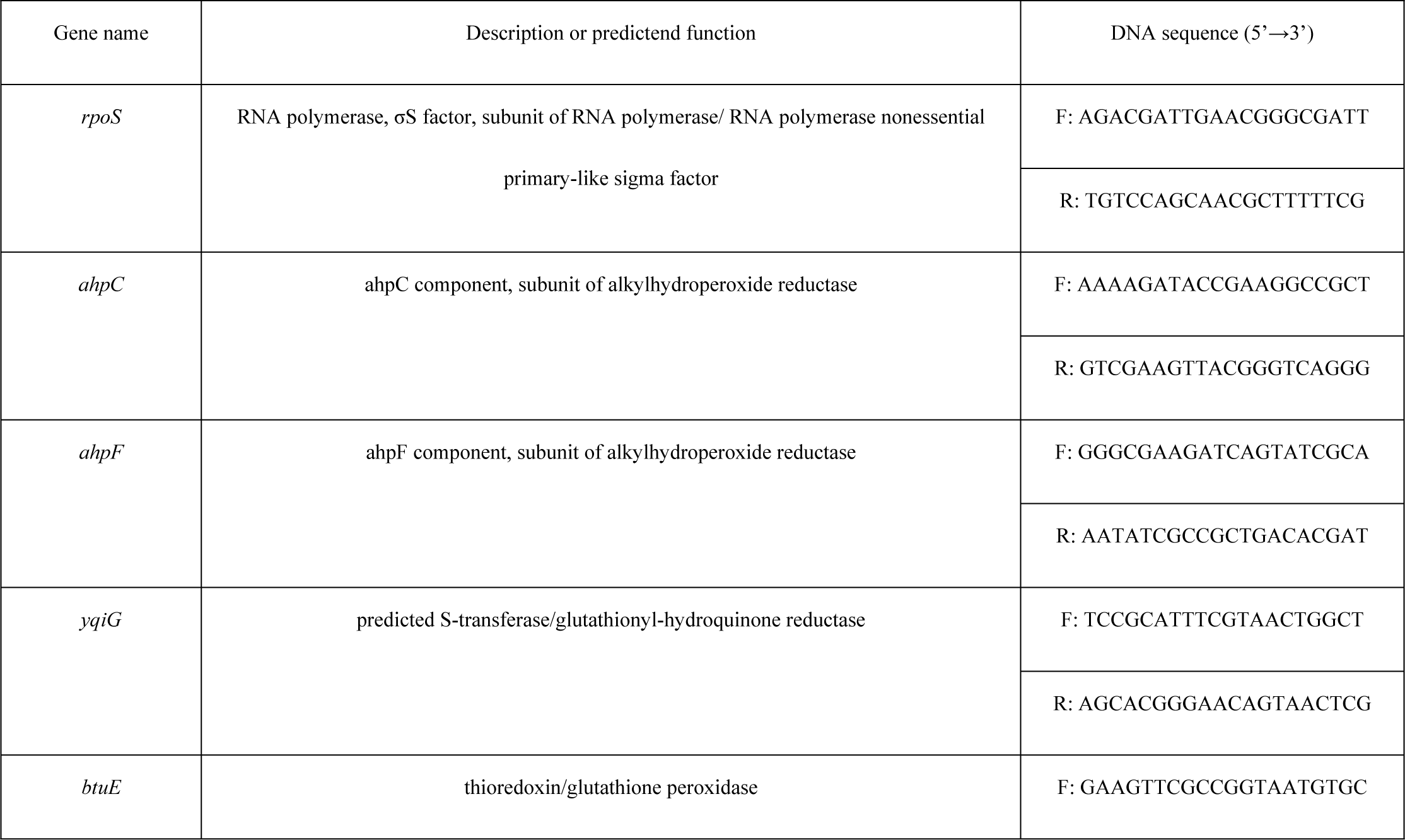
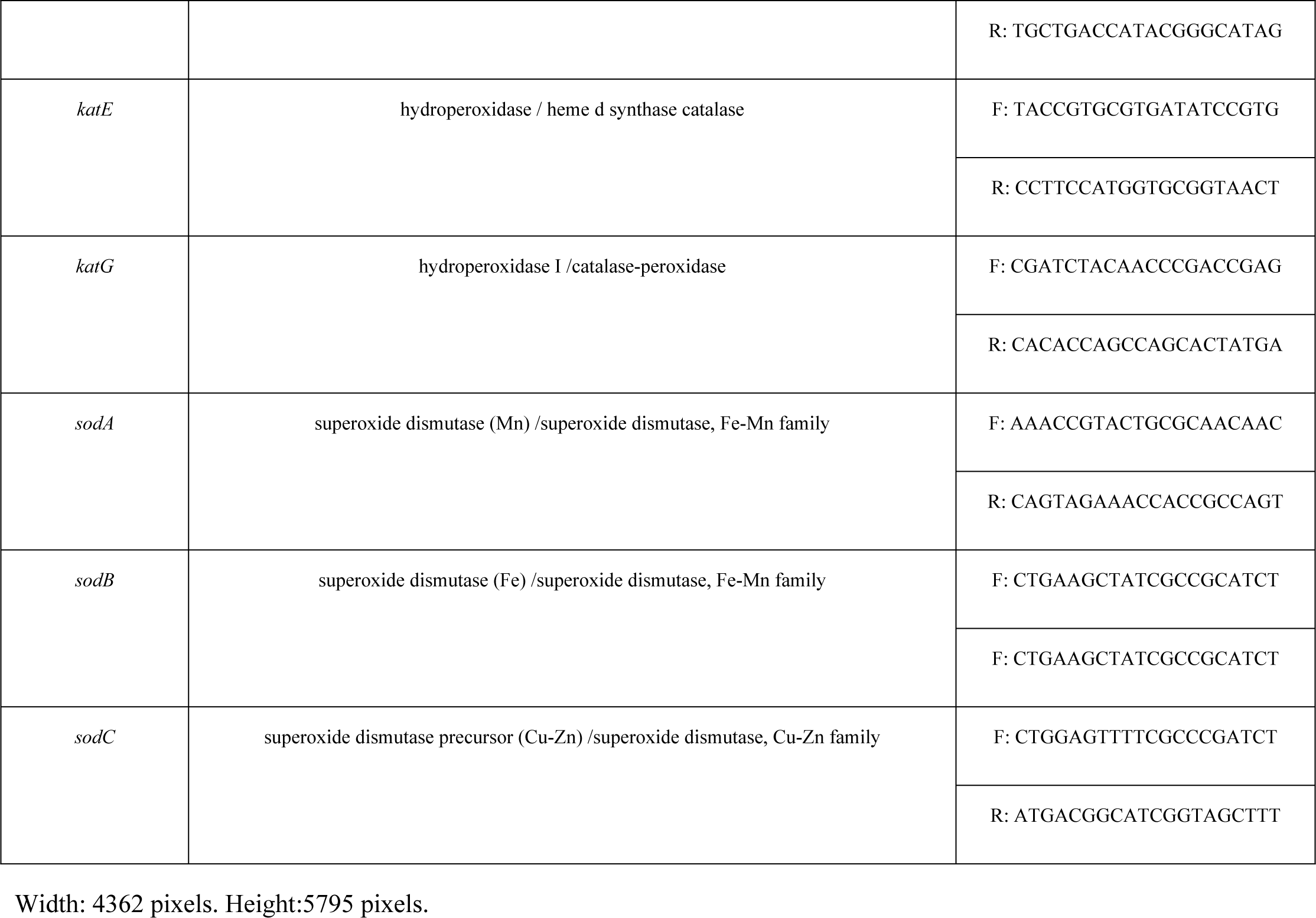
Primers used for the detection of hydrogen peroxide response genes during real-time PCR

## Results

### Construction and Identification of the *E. coli* BL21(DE3)^∆slp^

The *slp* gene mutant *E. coli* was constructed by the Targetron technique. Three positive mutant *E. coli* were selected. Genomic DNA of *E. coli* was extracted, and *slp* mutations were verified by PCR. As shown in Fig 1A, an approximately 1.5 kb gene fragment was amplified from the wild-type *E. coli* BL21(DE3) (Lane 1), and a 3.5 kb product was amplified from *E. coli* BL21(DE3)^∆slp^, which is 2kb larger than that in wild-type *E. coli* BL21(DE3), suggesting that the group II intron sequence was successfully inserted into the ORF (open reading frame) of the *slp* gene of *E. coli* BL21(DE3)^∆slp^ (Lanes 2-4). Sequencing revealed that the insertion site of the Group II intron was located at 161-162 bases in the open reading frame of the *slp* gene (data not shown). Real-time PCR was used to further examine the transcript expression of the *slp* gene in *E. coli*. The relative transcript expression of the *slp* gene in *E. coli* BL21(DE3)^∆slp^ was approximately 0.15-fold compared to the wild-type *E. coli* BL21(DE3) (Fig 1B). Therefore, *E. coli* BL21(DE3)^∆slp^ was successfully constructed.

**Fig 1.**
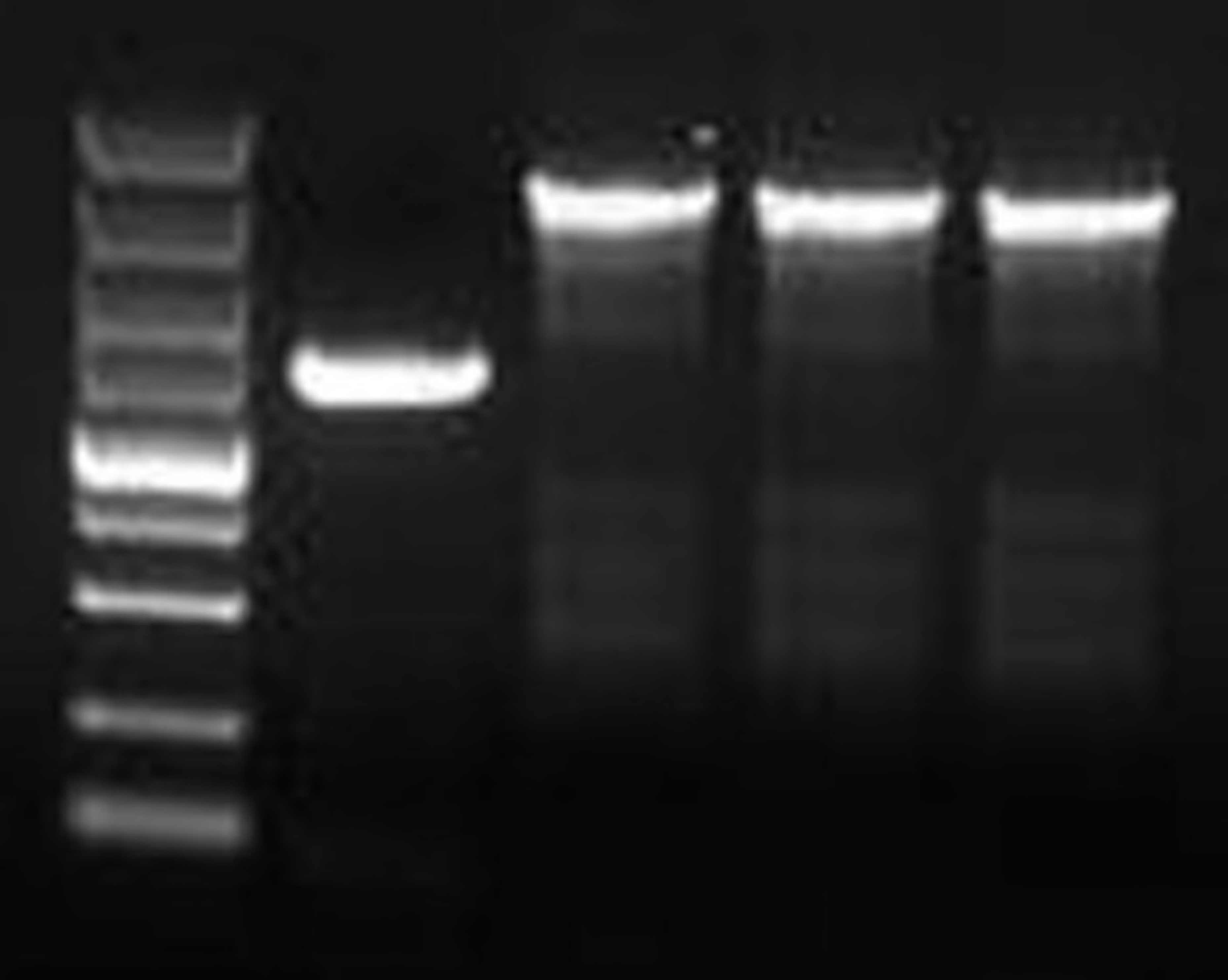
Identification of *Escherichia coli slp* gene mutation. **(A) Molecular weight changes of the *slp* gene in the genomic DNA**. Width: 2250 pixels (at 300 dpi). Height:1797 pixels (at 300 dpi). M: 5000 DNA marker; Lane 1: *slp* gene identification of wild-type *E. coli* BL21(DE3); Lanes 2-4: *slp* gene identification of *E. coli* BL21(DE3)^∆slp^. **(B) The relative transcript expression of the *slp* gene**. Width: 1889 pixels (at 600 dpi). Height:1686 pixels (at 600 dpi).

### The growth curves of *E. coli* under hydrogen peroxide stress

Cultured in LB liquid medium, there was no significant difference in the growth of *E. coli* BL21(DE3)^∆slp^ and wild-type *E. coli* BL21(DE3). Under 0.6% (v/v) H_2_O_2_ stress, the growths of *E. coli* BL21(DE3)^∆slp^ and wild-type *E. coli* BL21(DE3) were significantly inhibited (p<0.05), and *E. coli* BL21(DE3)^∆slp^ was significantly more sensitive than wild-type *E. coli* BL21(DE3) (p<0.05) (Fig 2). The result indicated that the mutation of the *slp* gene increased the sensitivity of *E. coli* under hydrogen peroxide stress.

**Fig 2.**
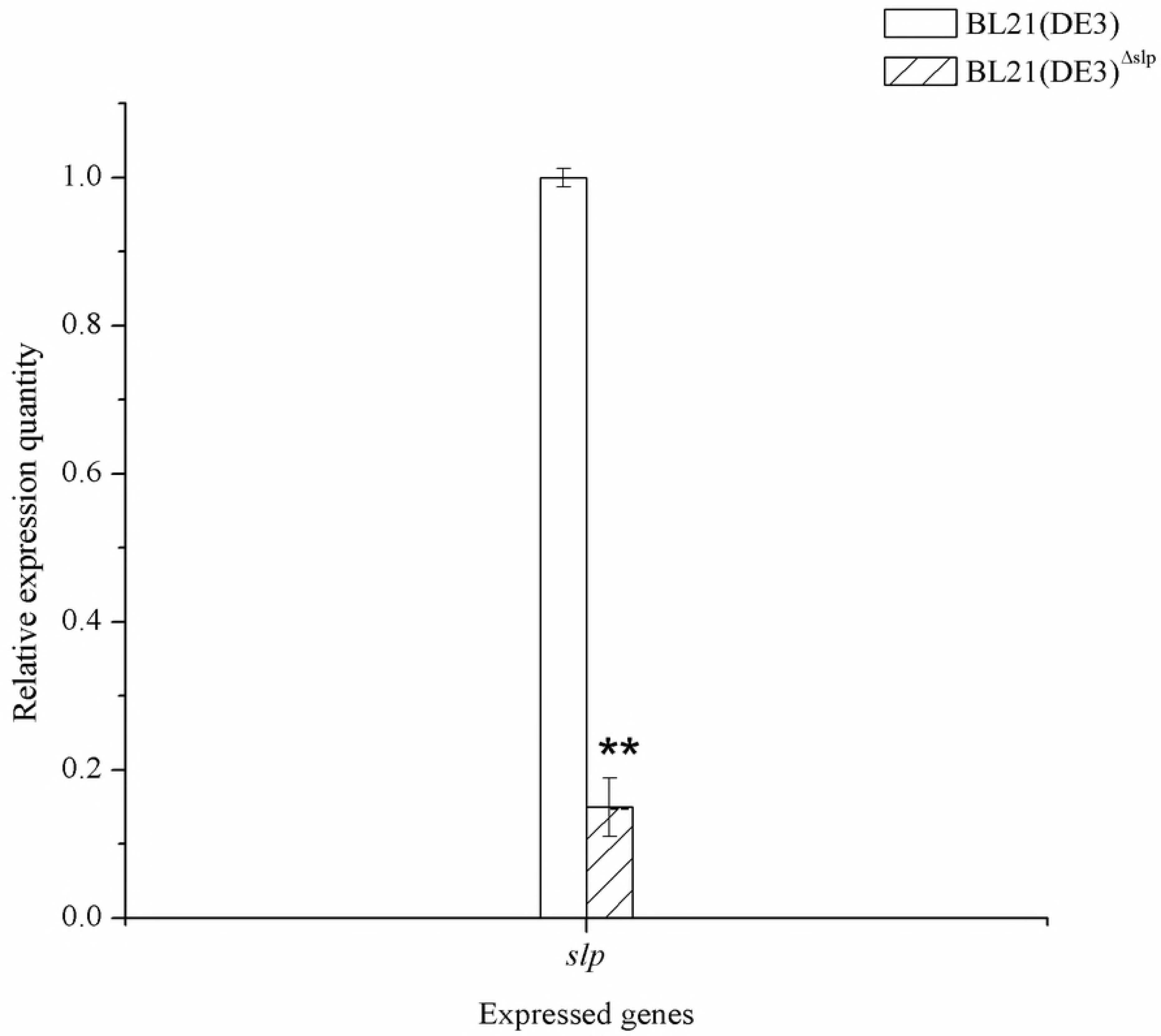
Growth curve of *E. coli* under hydrogen peroxide stress. Width: 1889 pixels (at 600 dpi). Height:1299 pixels (at 600 dpi).

### The output of RNA sequencing

Twelve transcriptome libraries were constructed with mRNA from *E. coli* BL21(DE3)^∆slp^ and wild-type *E. coli* BL21(DE3) in four groups, and three parallel libraries were included in each group. These libraries were sequenced using the Illumina HiSeqTM 2000 sequencing platform, and a total of 178,618,054 single raw reads were obtained. The read numbers in each library are shown in Table 3. After removing the non-coding RNA, the remaining clean reads in each library were mapped to wild-type *E. coli* BL21(DE3), and the mapping rates ranged from 86.52% to 90.43% (Table 3). The unique mapping rates from 12 libraries fluctuated within a relatively small range (from 86.52% to 89.67%, Table 3).

**Table 3.** The overview of RNA-Seq

| Treatment | Raw reads | Mapping reads | Mapping rate | Unique mapping |
| --- | --- | --- | --- | --- |
| C1- 1 | 15,520,272 | 14,034,449 | 90.43% | 89.67% |
| C1- 2 | 13,514,612 | 12,162,831 | 90.00% | 89.28% |
| C1- 3 | 15,227,790 | 13,666,726 | 89.75% | 89.06% |
| C2- 1 | 14,417,058 | 12,716,132 | 88.20% | 87.53% |
| C2- 2 | 14,026,030 | 12,398,647 | 88.40% | 87.73% |
| C2- 3 | 14,674,680 | 12,981,235 | 88.46% | 87.79% |
| C3- 1 | 15,069,934 | 13,451,699 | 89.26% | 88.72% |
| C3- 2 | 15,238,702 | 13,717,877 | 90.02% | 89.49% |
| C3- 3 | 15,245,556 | 13,713,991 | 89.95% | 89.41% |
| C4- 1 | 15,242,142 | 13,187,067 | 86.52% | 86.06% |
| C4- 2 | 15,334,084 | 13,354,249 | 87.09% | 86.62% |
| C4- 3 | 15,107,194 | 13,142,434 | 86.99% | 86.52% |

Width: 4362 pixels. Height:2043 pixels.

Wild-type *E. coli* BL21(DE3) of the C1 group grown in a non-hydrogen peroxide environment; *E. coli* BL21(DE3)^∆slp^ of the C2 group grown in a non-hydrogen peroxid e environment; Wild-type *E. coli* BL21(DE3) of the C3 group grown in 0.6% (vol/vol) H_2_O_2_; *E. coli* BL21(DE3)^∆slp^ of the C4 group grown in a 0.6% (vol/vol) H_2_O_2_. Three biological replicates per sample.

### Identification of the differentially expressed genes

To identify the gene expression signature induced by oxidative stress in *E. coli*, after the *slp* gene was mutated, the mutant and control strains were grown under 0.6% (vol/vol) hydrogen peroxide for three hours. Twelve samples passed our quality control criteria and were retained for further analyses (F test, univariate p< 0.05, FDR <0.001; Fig 3A). After filtering, we found 17,302 probes that were significantly deregulated at different time points, 1,879 that were upregulated and 1,625 downregulated (Fig 3B). Among all of these up/down-regulated genes, mutation of the *slp* gene directly caused the up-regulation of 13 genes (Fig 4A, C1 Vs C2) and down-regulation of 8 genes (Fig 4B, C1 Vs C2). In the hydrogen peroxide environment, the mutation of the *slp* gene caused an up-regulation of 63 genes (Fig 4A, C3 Vs C4). Hydrogen peroxide caused a significant up-regulation of 921 genes (Fig 4A, C1 Vs C3) and down-regulation of 839 genes (Fig 4B, C1 Vs C3) in wild-type *E. coli* BL21(DE3). In *E. coli* BL21(DE3)^∆slp^, hydrogen peroxide caused a significant up-regulation of 882 genes (Fig 4A, C2 Vs C4) and down-regulation of 778 genes (Fig 4B, C2 Vs C4).

**Fig 3.**
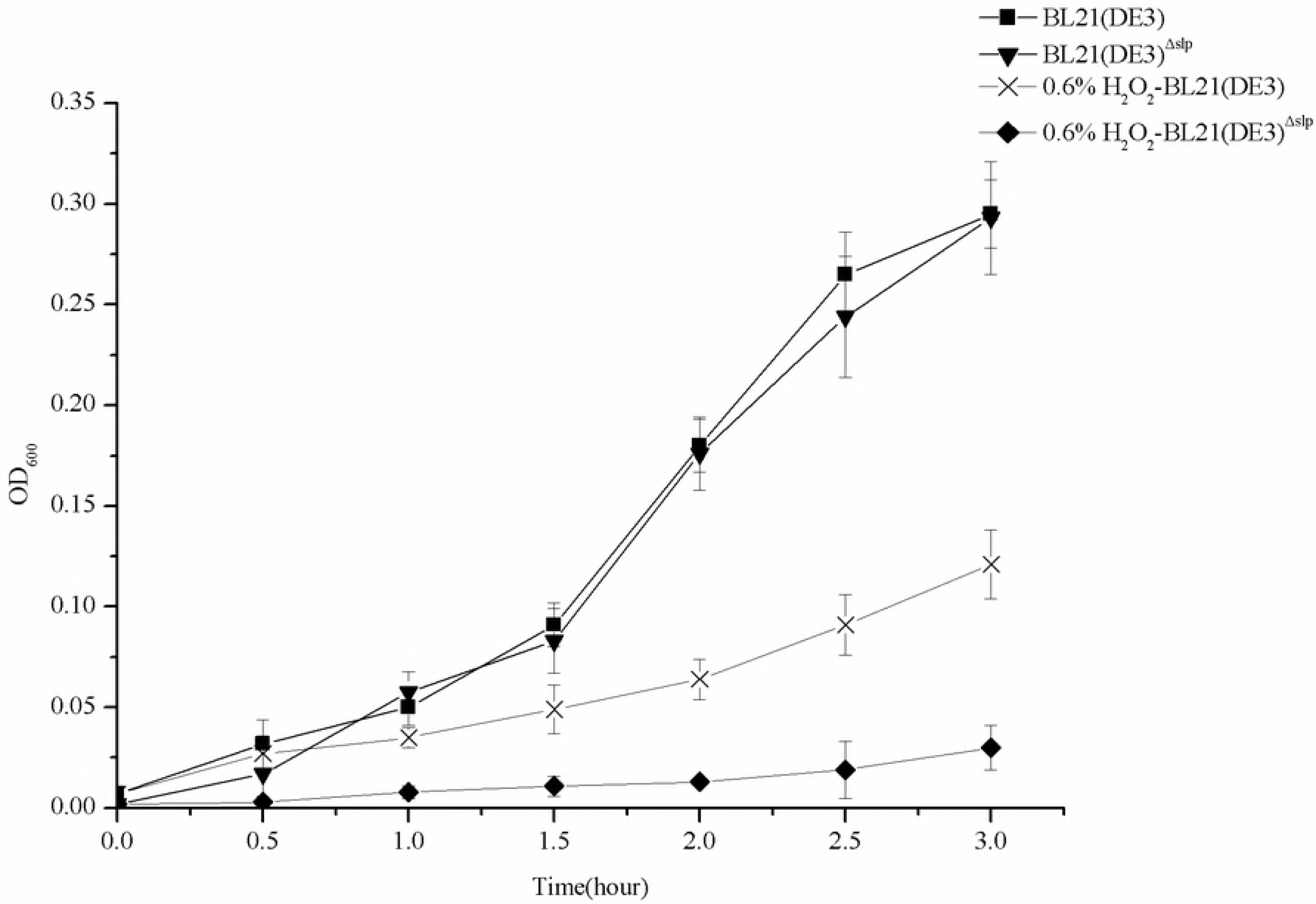
Differential gene analysis. **(A) Heatmap plot for the correlation coefficient between samples**. Width: 2250 pixels (at 300 dpi). Height:2250 pixels (at 300 dpi). The closer the linear relationship number of three biological samples is to 1, the higher the similarity between the parallel samples. Wild-type *E. coli* BL21(DE3) of the C1 group grown in a non-hydrogen peroxide environment; *E. coli* BL21(DE3)^∆slp^ of the C2 group grown in a non-hydrogen peroxide environment. Wild-type *E. coli* BL21(DE3) of the C3 group grown in a 0.6% (vol/vol) H_2_O_2_ environment. *E. coli* BL21(DE3)^∆slp^ of the C4 group grown in a 0.6% (vol/vol) H_2_O_2_ environment. (B) Visualization of differentially expressed genes. Width: 1181 pixels (at 300 dpi). Height:1181 pixels (at 300 dpi). -genes that were significantly up-regulated. -genes that were significantly down-regulated. -genes that were not significantly changed. a: Differentially expressed genes that were significantly altered after mutation of the *slp* gene. b: Differentially expressed genes caused by 0.6% (vol/vol) H_2_O_2_ environment in wild-type *E. coli* BL21(DE3). c: Differentially expressed genes caused by 0.6% (vol/vol) H_2_O_2_ in *E. coli* BL21(DE3)^∆slp^. d: Differentially expressed genes that were significantly altered after mutation of the *slp* gene under 0.6% (vol/vol) H_2_O_2_.

**Fig 4.**
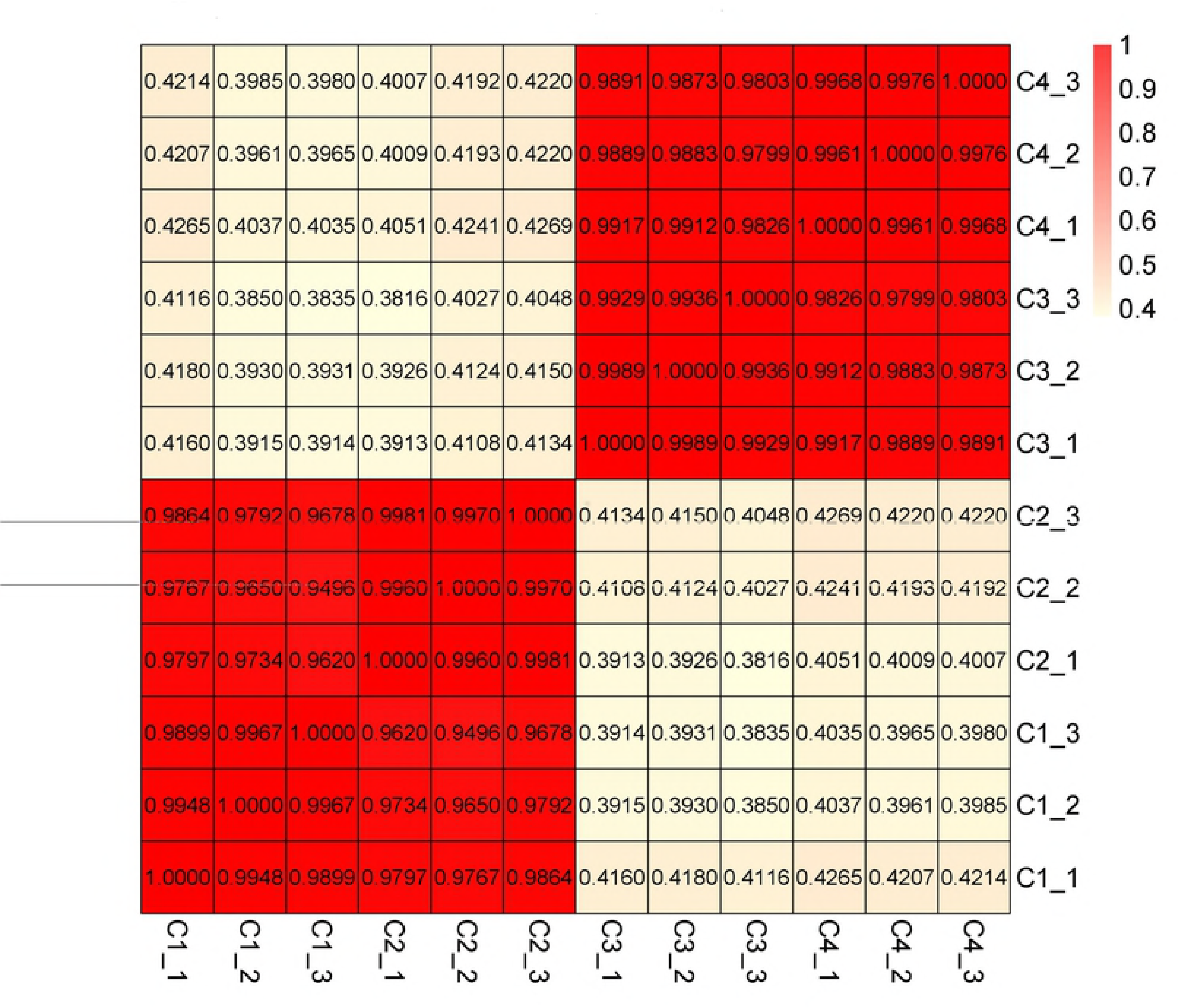
Venn map of the up/down-regulated genes. **(A) Up-regulated genes**. Width: 2250 pixels (at 300 dpi). Height:2250 pixels (at 300 dpi). (B) Down-regulated genes. Width: 2250 pixels (at 300 dpi). Height:2250 pixels (at 300 dpi). Wild-type *E. coli* BL21(DE3) of the C1 group grown in a non-hydrogen peroxide environment; *E. coli* BL21(DE3)^∆slp^ of the C2 group grown in a non-hydrogen peroxide environment; Wild-type *E. coli* BL21(DE3) of the C3 group grown in 0.6% (vol/vol) H_2_O_2_; *E. coli* BL21(DE3)^∆slp^ of the C4 group grown in 0.6% (vol/vol) H_2_O_2_.

### The differentially expressed genes related to the hydrogen peroxide oxidative stress pathway

Significant changes in oxidative stress, glucose metabolism, and energy metabolism pathways were observed after the *slp* gene was mutated (Table 4). After the *slp* gene was mutated, it mainly resulted in a significant increase in the genes in the glucose metabolism pathway (GO: 0004641; GO: 0004637; GO: 0004644; GO: 0009401) (Table 4). In the energy metabolism pathway, it caused a significant decrease in the gene (GO: 0015419; GO: 0005524); however, there was dramatically increased expression of the genes in the energy metabolism pathway when *E. coli* BL21(DE3)^∆slp^ was grown in a 0.6% (vol/vol) hydrogen peroxide environment (Table 4).

**Table 4.** The major significantly represented GO terms of the genes

| Pathway | GO id | GO description |  | log2Fold<br>Change | q value | Up/Down-Regulation |
| --- | --- | --- | --- | --- | --- | --- |
| Oxidative stress | GO:0004364 | glutathione transferase activity | C1 Vs C2 | -1.0148 | 8.28E-202 | Down |
|  |  |  | C2 Vs C4 | 2.7097 | 7.65E-164 | Up |
|  |  |  | C1 Vs C3 | 1.0146 | 8.28E-202 | * |
|  |  |  | C3 Vs C4 | - | - | - |
|  | GO:0004602 | glutathione peroxidase activity | C1 Vs C2 | 2.2073 | 1.42E-38 | Up |
|  |  |  | C2 Vs C4 | 1.003 | 2.52E-37 | * |
|  |  |  | C1 Vs C3 | 1.0288 | 1.42E-38 | * |
|  |  |  | C3 Vs C4 | - | - | - |
|  | GO:0004096 | catalase activity | C1 Vs C2 | 2.8124 | 1.80E-238 | Up |
|  |  |  | C2 Vs C4 | 3.9789 | 0 | Up |
|  |  |  | C1 Vs C3 | 1.2577 | 1.80E-238 | Up |
|  |  |  | C3 Vs C4 | - | - | - |
|  | GO:0055114 | oxidation-reduction process | C1 Vs C2 | 1.1566 | 4.18E-07 | Up |
|  |  |  | C2 Vs C4 | -3.9447 | 5.65E-296 | Down |
|  |  |  | C1 Vs C3 | -2.5204 | 1.57E-18 | Down |
|  |  |  | C3 Vs C4 | - | - | - |
| Sugar metabolism | GO:0004641 | phosphoribosyl<br>formylglycinamide cyclo-ligase<br>activity | C1 Vs C2 | 1.0378 | 6.81E-28 | Up |
|  |  |  | C2 Vs C4 | - | - | - |
|  |  |  | C1 Vs C3 | - | - | - |
|  |  |  | C3 Vs C4 | - | - | - |
|  | GO:0004637 | phosphoribosylamine-glycine<br>ligase activity | C1 Vs C2 | 1.3473 | 1.27E-31 | Up |
|  |  |  | C2 Vs C4 | -2.8232 | 8.34E-163 | Down |
|  |  |  | C1 Vs C3 | -1.6542 | 4.02E-31 | Down |
|  |  |  | C3 Vs C4 | - | - | - |
|  | GO:0004644 | phosphoribosylg lycinamide<br>formyltransferase activity | C1 Vs C2 | 1.1033 | 4.25E-26 | Up |
|  |  |  | C2 Vs C4 | - | - | - |
|  |  |  | C1 Vs C3 | - | - | - |
|  |  |  | C3 Vs C4 | - | - | - |
|  | GO:0009401 | phosphoenolpyruvate-dependent<br>sugar phosphotransferase system | C1 Vs C2 | 1.2398 | 1.58E-18 | Up |
|  |  |  | C2 Vs C4 | - | - | - |
|  |  |  | C1 Vs C3 | - | - | - |
|  |  |  | C3 Vs C4 | - | - | - |
| Energy metabolism | GO:0015419 | sulfate<br>transmembrane-transporting<br>ATPase activity | C1 Vs C2 | -1.0169 | 2.21E-15 | Down |
|  |  |  | C2 Vs C4 | 3.9955 | 2.98E-289 | Up |
|  |  |  | C1 Vs C3 | 1.8282 | 5.36E-79 | Up |
|  |  |  | C3 Vs C4 | - | - | - |
|  | GO:0005524 | ATP binding | C1 Vs C2 | -1.0825 | 1.09E-05 | Down |
|  |  |  | C2 Vs C4 | 4.8281 | 8.14E-71 | Up |
|  |  |  | C1 Vs C3 | 2.4912 | 1.50E-16 | Up |
|  |  |  | C3 Vs C4 | - | - | - |

\* represents no significant change; - represents no detected change. Wild-type *E. coli* BL21(DE3) of the C1 group grown in a non-hydrogen peroxide environment; *E. coli* BL21(DE3)<sup>Δslp</sup> of the C2 group grown in a non-hydrogen peroxide environment; Wild-type *E. coli* BL21(DE3) of the C3 group grown in 0.6% (vol/vol) H<sub>2</sub>O<sub>2</sub>; *E. coli* BL21(DE3)<sup>Δslp</sup> of the C4 group grown in 0.6% (vol/vol) H<sub>2</sub>O<sub>2</sub>.

* represents no significant change; - represents no detected change. Wild-type *E. coli* BL21(DE3) of the C1 group grown in a non-hydrogen peroxide environment; *E. coli* BL21(DE3)^∆slp^ of the C2 group grown in a non-hydrogen peroxide environment; Wild-type *E. coli* BL21(DE3) of the C3 group grown in 0.6% (vol/vol) H_2_O_2_; *E. coli* BL21(DE3)^∆slp^ of the C4 group grown in 0.6% (vol/vol) H_2_O_2_.

In the oxidative stress pathway, the change in antioxidant enzymes was the main concern. When wild-type *E. coli* BL21(DE3) was exposed to a 0.6% (vol/vol) hydrogen peroxide environment, the gene encoding glutathione transferase (GO:0004364) or glutathione peroxidase (GO:0004602) was not changed, and the gene encoding catalase (GO:0004096) was increased approximately 1.26-fold (Table 4). Catalase (approximately 2.81-fold) and glutathione peroxidase (approximately 2.21-fold) were significantly increased after *slp* gene mutation, while glutathione transferase decreased (approximately 1.01-fold) (Table 4). When *E. coli* BL21(DE3)^∆slp^ was cultured in 0.6% (vol/vol) hydrogen peroxide, the expression of genes encoding catalase (approximately 3.98-fold) and glutathione transferase (approximately 2.71-fold) were dramatically increased, while the glutathione peroxidase gene had normal expression (Table 4). These antioxidant enzymes were characterized by the significantly represented GO terms of *E. coli* BL21(DE3)^∆slp^, which were responsive to 0.6% (vol/vol) H_2_O_2_ oxidative stress.

### Enzyme activity

We further examined the oxidative stress-related activities. As shown in Fig 5, catalase activities in *E. coli* BL21(DE3)^∆slp^ were significantly higher (p<0.05) than in wild-type *E. coli* BL21(DE3) during 3 hours of culture, which suggested that the mutation of the *slp* gene caused a marked increase in the catalase activity in *Escherichia coli*. Under 0.6% (vol/vol) H_2_O_2_ oxidative stress, the catalase activities of *E. coli* BL21(DE3)^∆slp^ and wild-type *E. coli* BL21(DE3) were significantly increased (p<0.05), and the catalase activities in *E. coli* BL21(DE3)^∆slp^ were obviously higher than those in the wild-type strain from 1 to 3 hours (p <0.05).

**Fig 5.**
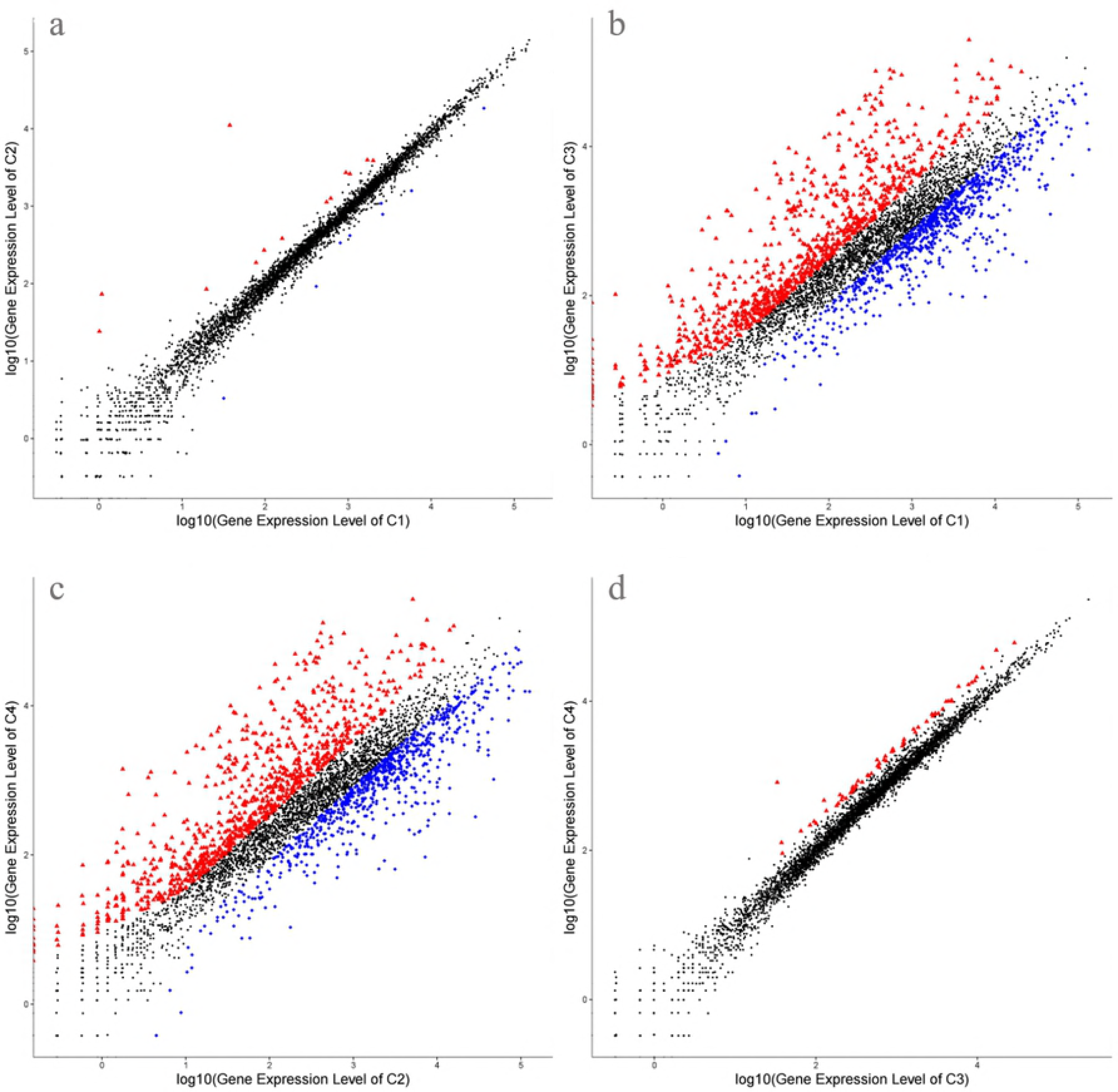
Catalase activity of *E. coli* under hydrogen peroxide stress. Width: 1889 pixels (at 600 dpi). Height:1516 pixels (at 600 dpi).

The glutathione enzyme activity of *E. coli* is shown in Fig 6A, and the glutathione S-transferase activities in *E. coli* BL21(DE3)^∆slp^ were markedly lower (p<0.05) than that in wild-type *E. coli* BL21(DE3), which suggested that the mutation of the *slp* gene caused a marked decrease in the glutathione S-transferase activity in *Escherichia coli*. Under 0.6% (vol/vol) H_2_O_2_ oxidative stress, no change was found in *E. coli* BL21(DE3)^∆slp^ and wild-type *E. coli* BL21(DE3). Moreover, the mutation of the *slp* gene caused a significant increase in glutathione peroxidase activity in *E. coli* compared to the control. Additionally, 0.6% (vol/vol) H_2_O_2_ addition did not influence the activity of glutathione peroxidase both in *E. coli* BL21(DE3)^∆slp^ or wild-type *E. coli* BL21(DE3) (Fig 6B).

**Fig 6.**
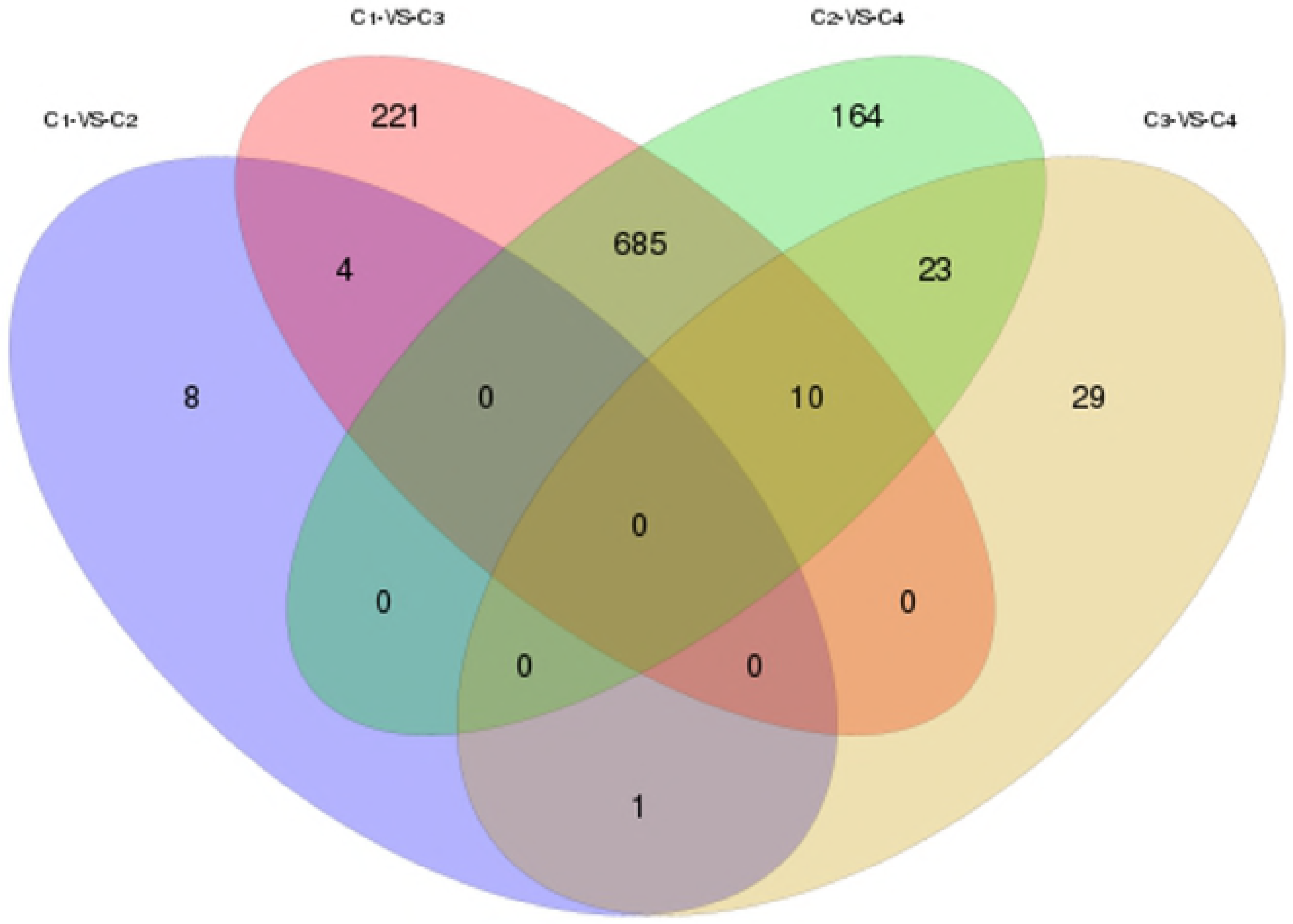
Glutathione activity of *E. coli* under hydrogen peroxide stress. **(A) Glutathione S-transferase activity**. Width: 1889 pixels (at 600 dpi). Height:1410 pixels (at 600 dpi). **(B) Glutathione peroxidase activity**. Width: 1889 pixels (at 600 dpi). Height:1410 pixels (at 600 dpi).

### Trace Malondialdehyde contents in *E. coli*

As shown in Fig 7, mutations of the *slp* gene in wild-type *E. coli* BL21(DE3) had no significant effect on the content of trace malondialdehyde during 3 hours of culture compared with the control. The content of trace malondialdehyde in *E. coli* BL21(DE3)^∆slp^ and wild-type *E. coli* BL21(DE3) significantly increased from 0.5 hours to 2 hours under 0.6% (vol/vol) H_2_O_2_ oxidative stress (p<0.05); however, the content of malondialdehyde in *E. coli* BL21(DE3)^∆slp^ was significantly higher between 1.5 hours and 2 hours than wild-type *E. coli* BL21(DE3) (p < 0.05).

**Fig 7.**
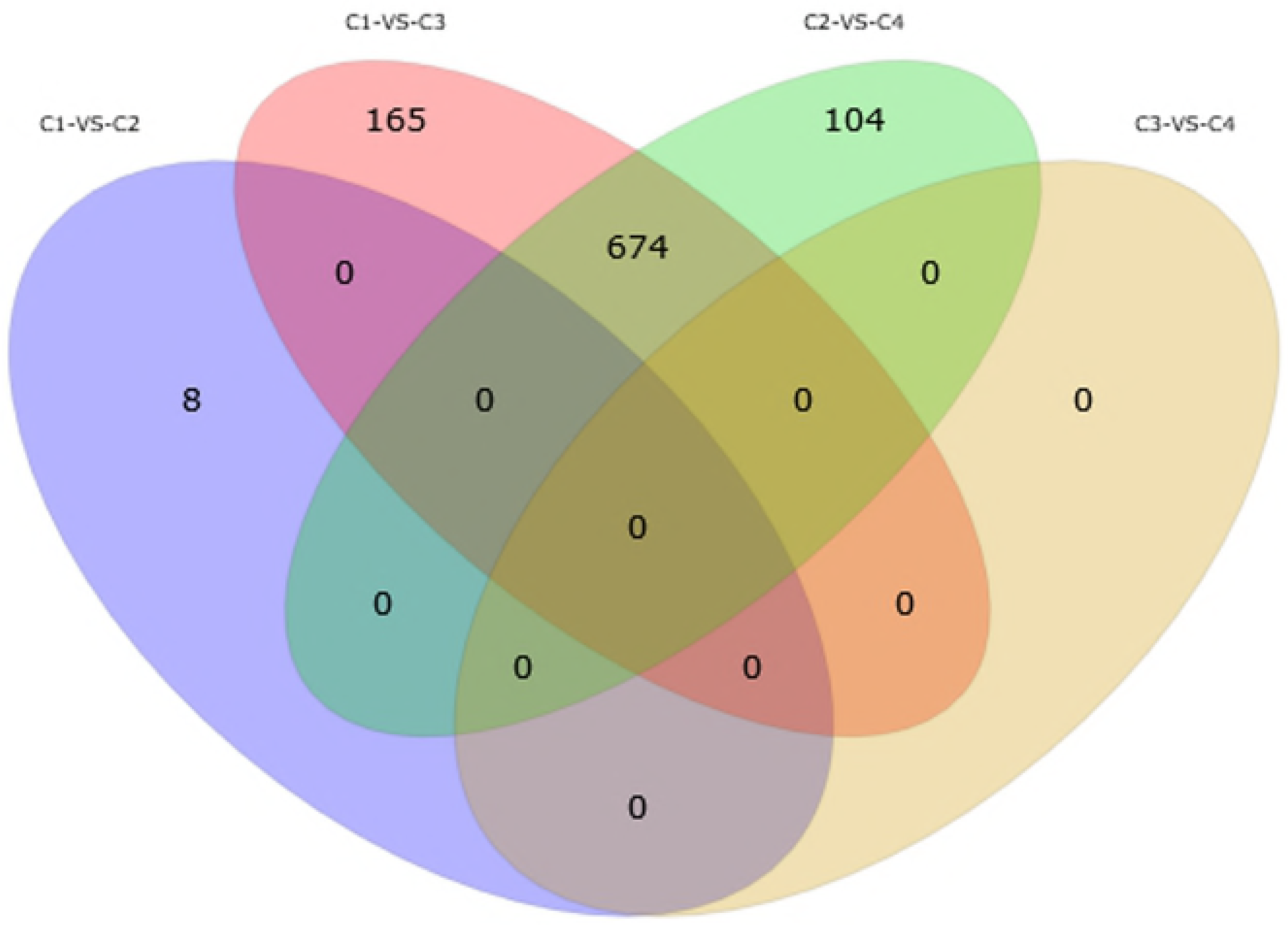
Trace malondialdehyde content of *E. coli* under hydrogen peroxide stress. Width: 1889 pixels (at 600 dpi). Height:1471 pixels (at 600 dpi).

### Transcript expression of antioxidase-related genes

Among all of the detected genes, as shown in Fig 8, after the *slp* gene mutation, there was no significant change in the *katE* gene encoding the HPII family and the *ahpCF* gene encoding alkyl peroxide reductase AhpCF, as well as in the expression of the *RpoS* (encoding σ^S^ regulator) (P<0.05). But the expression of the genes encoding glutathione (*yqjG*, *btuE*) and superoxide dismutase (*sodB*, *sodC*) increased remarkably. Simultaneously, the transcript expression of *katG* (encoding HPI family) also significantly increased (P<0.05). When wild-type *E. coli* BL21(DE3) was grown in 0.6% (v/v) H_2_O_2_ oxidative stress, the *sodB* gene was significantly down-regulated, and the expression of the *RpoS* gene was not changed, while all of the other genes were significantly increased (P<0.05). With the exception of the *sodB* gene, which was significantly down-regulated, all other genes were remarkedly increased when *E. coli* BL21(DE3)^∆slp^ was grown in 0.6% (vol/vol) H_2_O_2_ oxidative stress. The expression levels of the *katG* and *butE* genes were increased in both *E. coli* BL21(DE3)^∆slp^ and wild-type *E. coli* BL21(DE3) under 0.6% (vol/vol) H_2_O_2_ oxidative stress, but the expression of these two genes increased more in the mutant. Additionally, the expression of the *katE* gene in wild-type *E. coli* BL21(DE3) increased more obviously.

**Fig 8.**
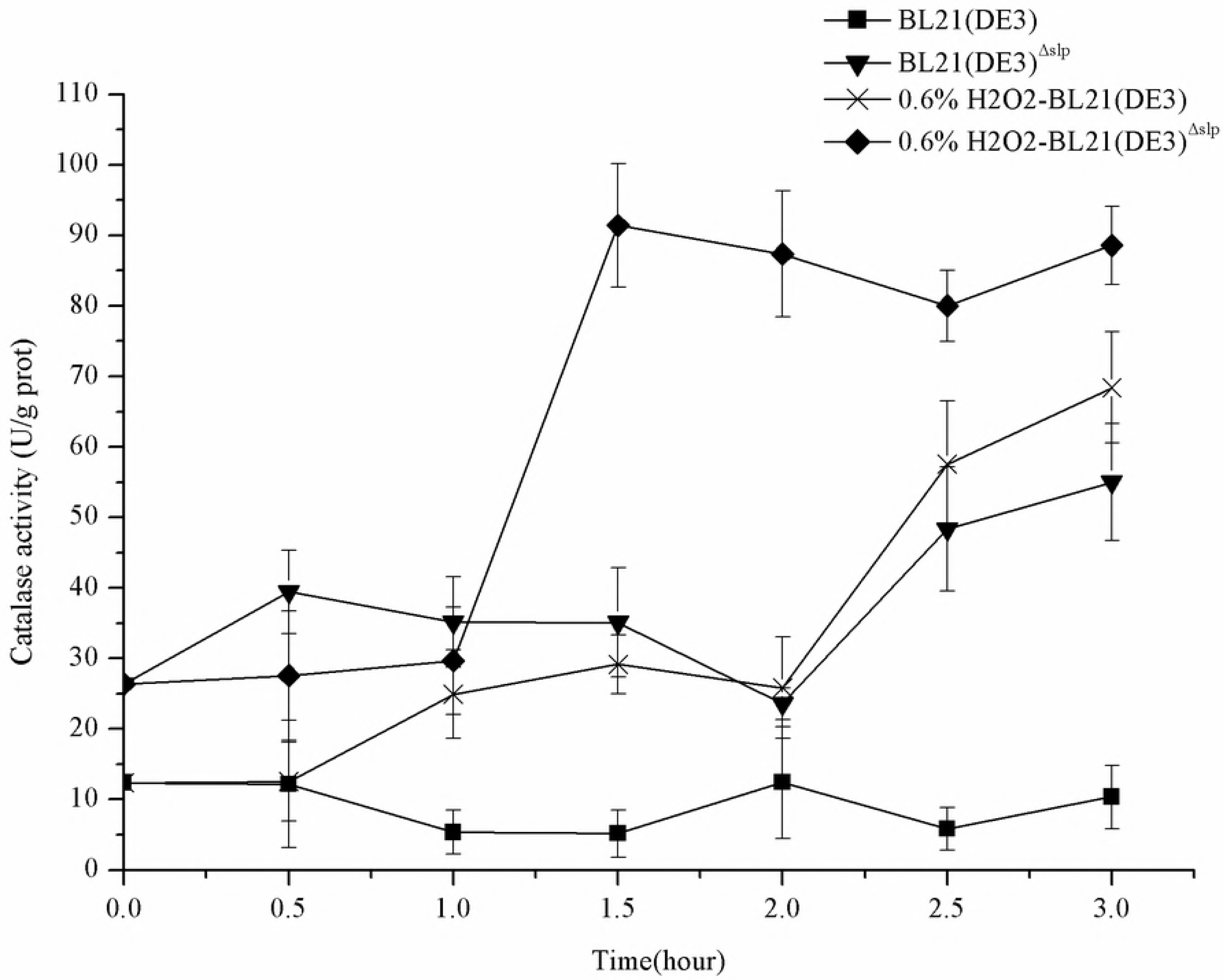
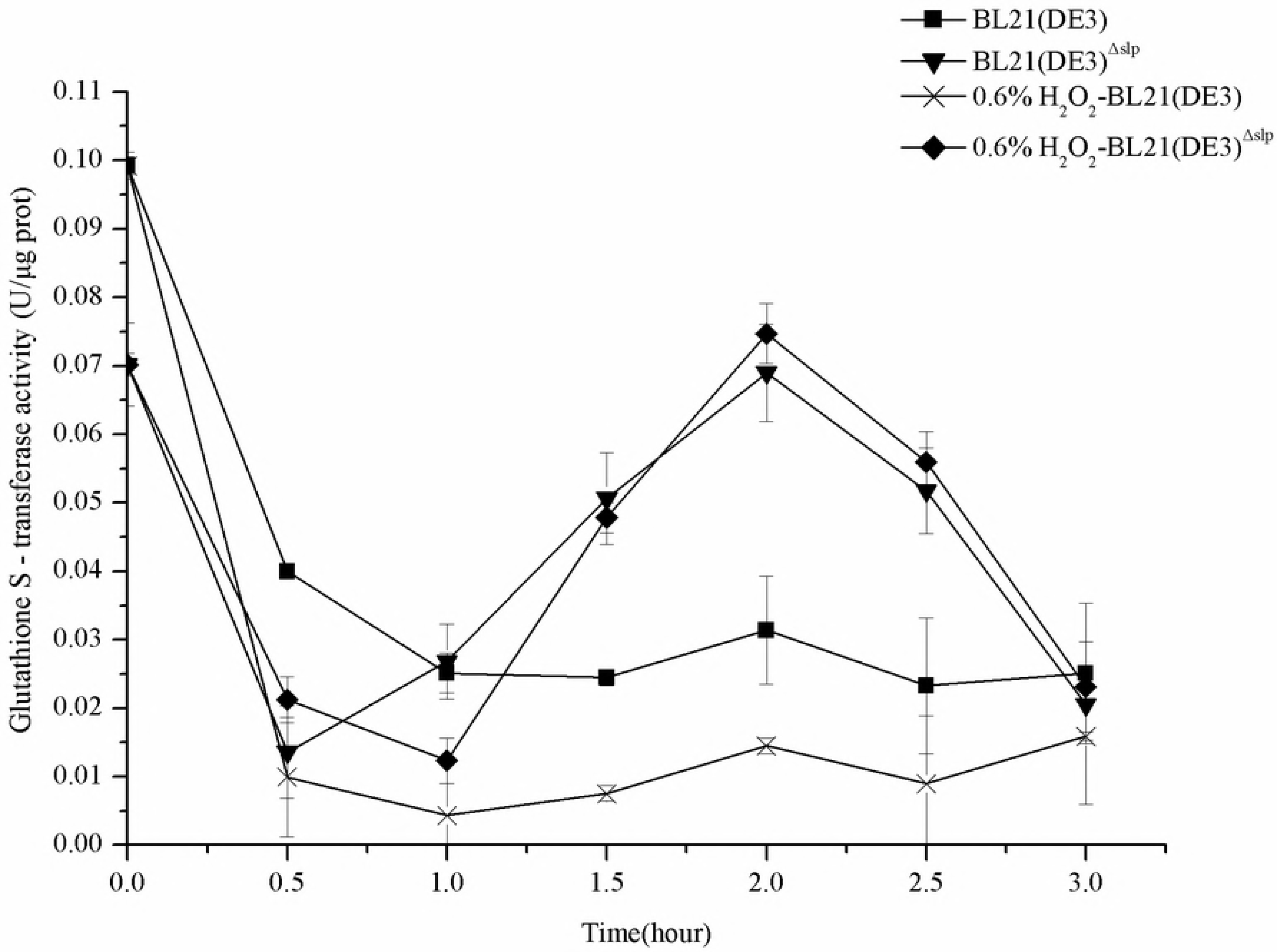
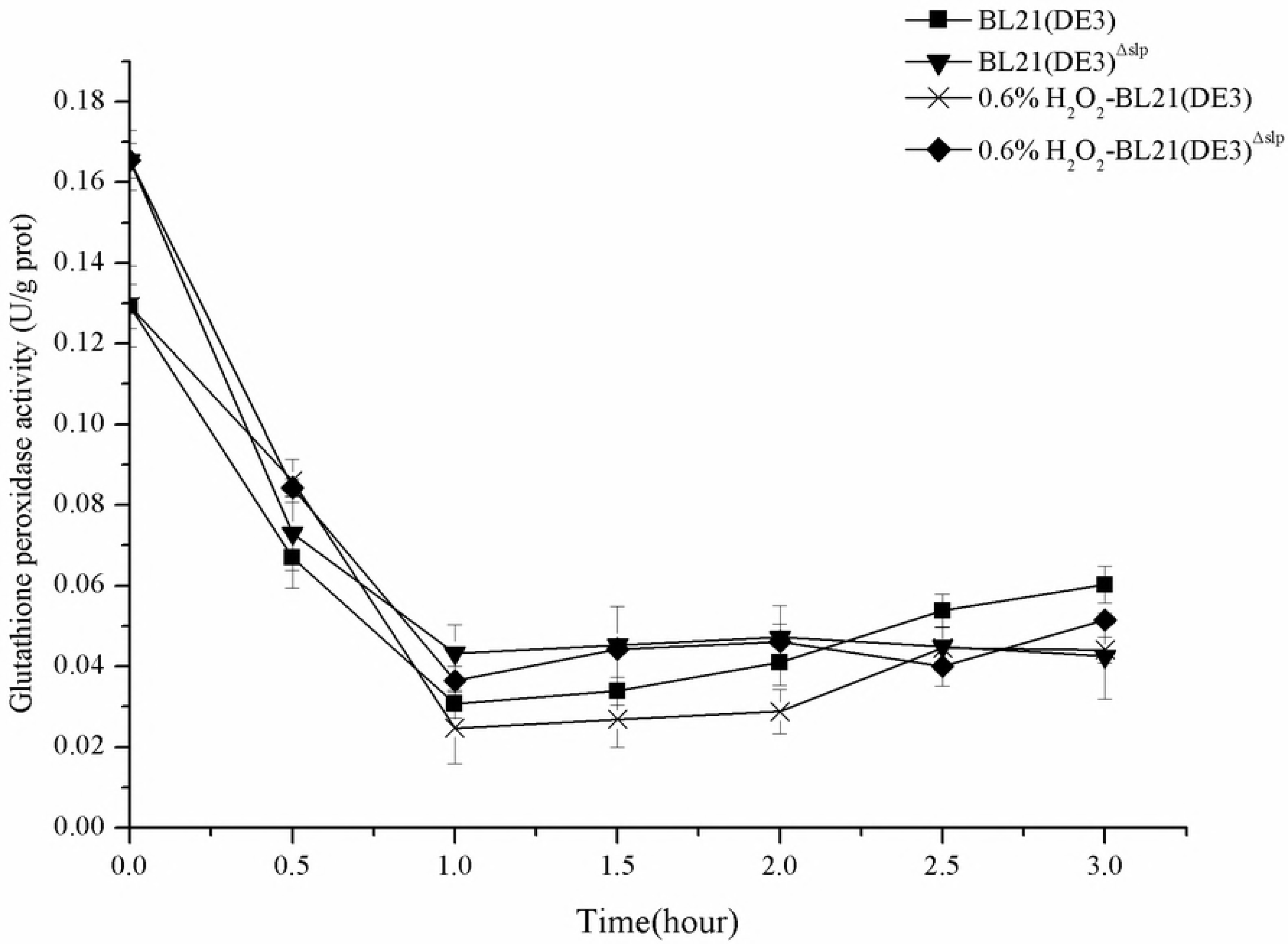
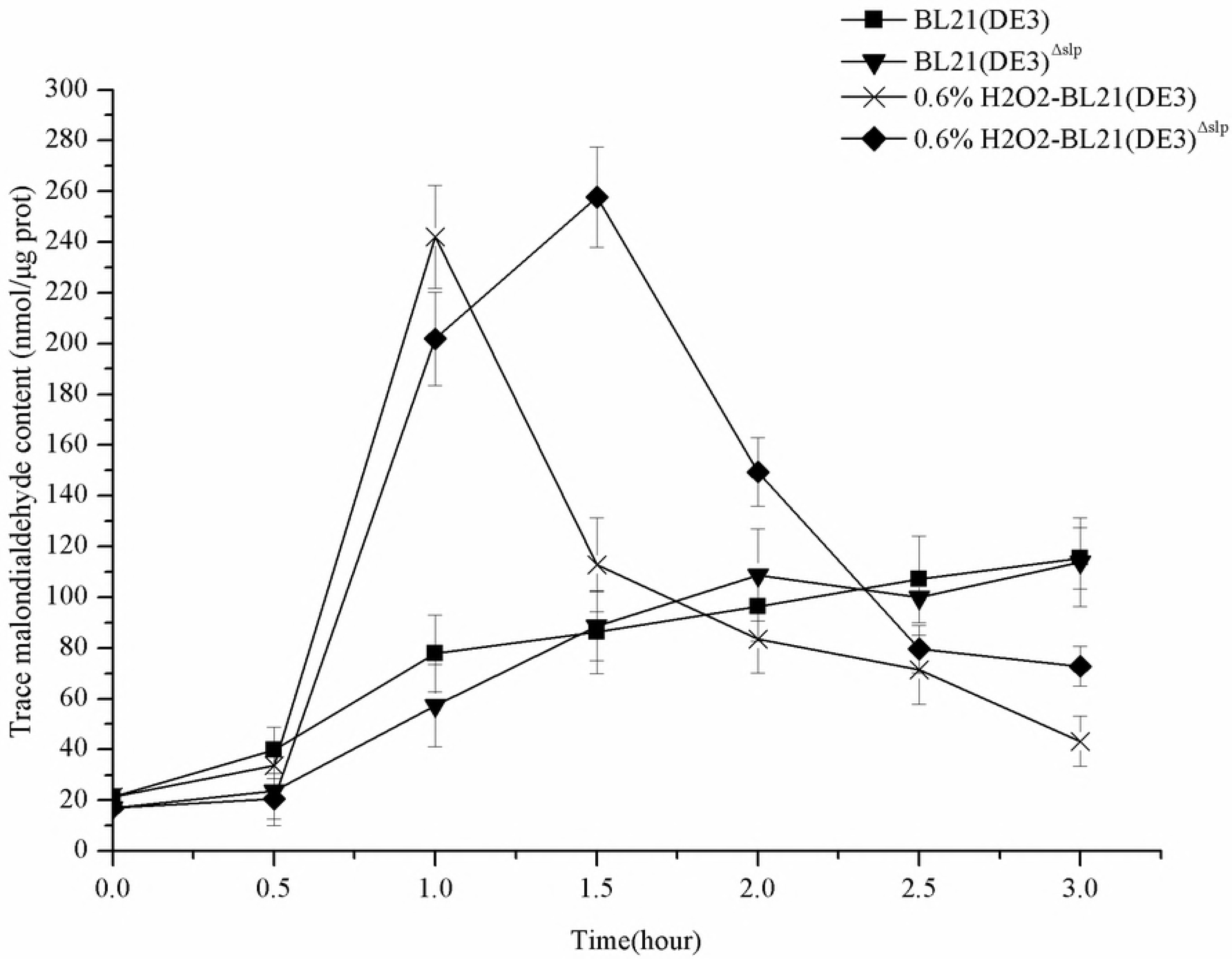
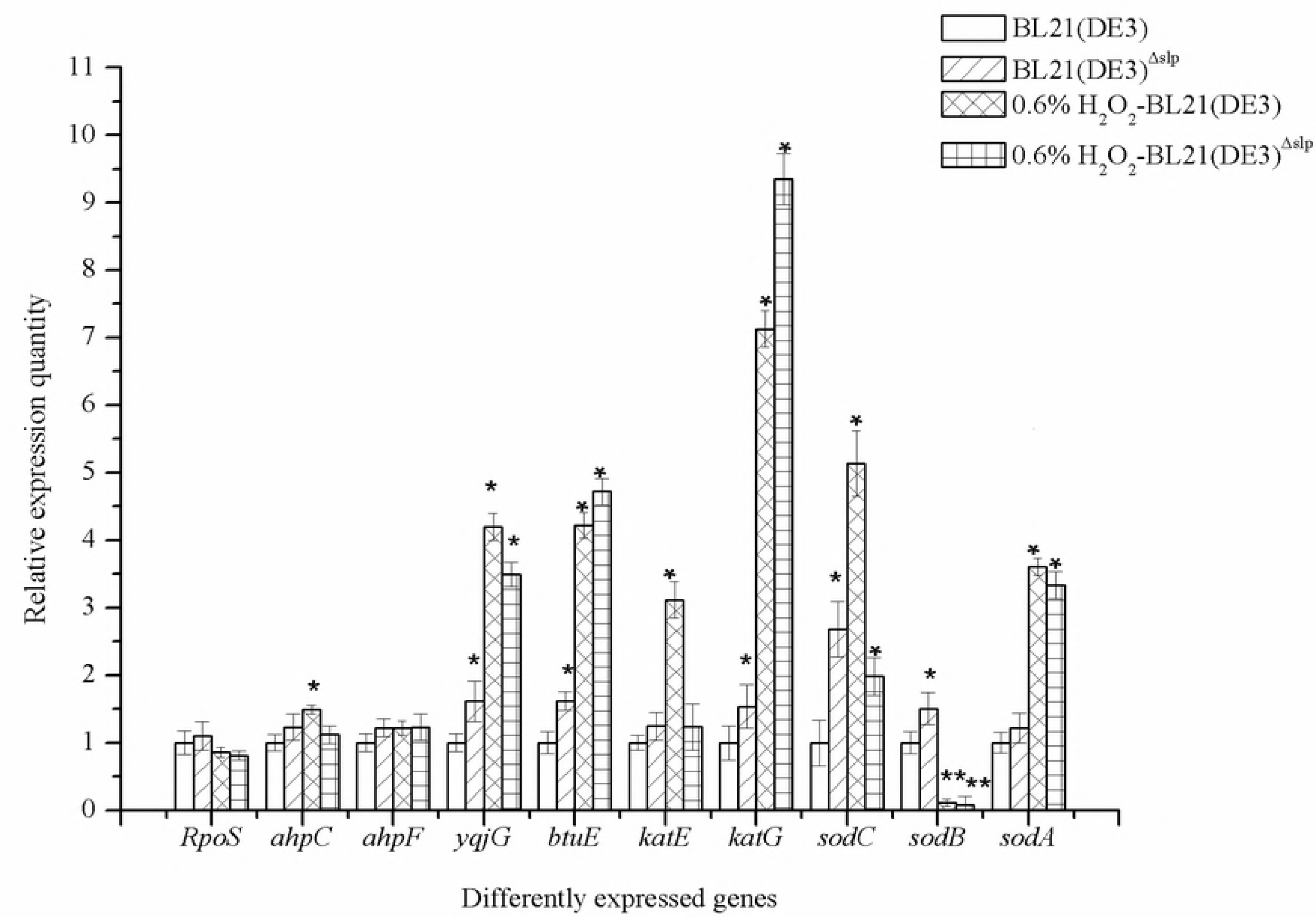
Expression of antioxidant-related genes in *E. coli* under hydrogen peroxide stress. Width: 1889 pixels (at 600 dpi). Height:1334 pixels (at 600 dpi).

## Discussion

The localization of carbon starvation-induced lipoprotein Slp in the outer membrane suggests a potential role in protecting stationary phase cells from environmental stress or facilitating nutrient availability in the periplasm [9]. A previous study suggested that it could be induced by carbon deficiency or oxidative stress [4]. However, there was no clear conclusion about the oxidative stress induction mechanism of carbon starvation-induced lipoprotein Slp.

In this study, we first constructed an *slp* null mutant *E. coli* using the Group II intron insertion method (Fig 1). Group II introns use a remarkable mobility mechanism in which the excised intron RNA uses its ribozyme activity to insert directly into a DNA target site by reverse-splicing, termed retro-homing [22]. Most of the target specificity comes from the base-pairing of the intron RNA to the DNA target sequence, which means that it is possible to reprogram group II introns to insert into desired sites simply by modifying the intron RNA [23]. Group II introns are minimally dependent on host factors and widely applicable to a wide variety of bacteria. Mutants have been successfully constructed in various strains, such as *E. coli*, *S. typhimurium* and *L. lactis* [23, 24].

We discovered that the *slp* gene mutant strain had increased susceptibility to 0.6% (vol/vol) H_2_O_2_ oxidative stress and grew poorly with oxygen during the logarithmic phase (Fig 2). Gregory found that there are no significant changes in resistance to oxidative stress by measuring zones of inhibition surrounding disks containing 10 µl of 50% peroxide in the *slp* gene mutant strain; however, this study was not designed to measure slight differences in the sensitivity of the *slp* gene mutant to stress treatments [9]. During entry into the stationary phase, *E. coli* including *E. coli* BL21(DE3)^∆slp^ devoted a considerable amount of synthetic protein to resist adverse environments during long-term exposure to hydrogen peroxide [2]. After *slp::phoA* strains were subjected to oxidative challenge and followed for 4 hours of glucose starvation during exponential growth, Alexander indicated that alteration of Slp did not change starvation-induced cross resistance to osmotic, thermal or oxidative stress [4]. Unfortunately, Alexander studied whether *slp::phoA* strains could depend on Pex protein protection and thus produce starvation-induced cross-resistance. No studies confirmed that carbon starvation induced-lipoprotein Slp is dependent on cAMP/CRP, Pex or cross-protection.

The function of Slp protein was predicted at the transcript level in *E. coli*, and it was found that the mutation of the *slp* gene caused a change in the oxidative stress pathway, among which the expression of antioxidant enzyme genes was the most obvious, indicating that the Slp protein had an effect on the antioxidant process (Table 4). In addition, the function of the Slp protein may also involve sugar metabolism and energy metabolism pathways (Table 4). Membrane lipoproteins in *E. coli* play an important physiological role in signal transduction, substance transport and so on [25]. In the hydrogen peroxide environment, mutations of the *slp* gene caused increased expression of iron-related genes (data not shown), and we speculate that the Slp protein has a similar function as the Dps protein and could chelate free iron to reduce the damage of hydrogen peroxide on cells [26]. Free iron ions could promote the Fenton reaction of hydrogen peroxide, causing serious production of proteins, nucleic acids, and lipid molecules [27].

Indeed, H_2_O_2_ itself can potentially damage enzymes by oxidizing sulfhydryl and iron-sulfur moieties. Upon conversion to a hydroxyl radical, it produces mutagenic and lethal lesions [28]. Catalase in the body uses iron porphyrin as the auxiliary base, and catalytic decomposition of H_2_O_2_ and removal from the body of H_2_O_2_ prevents cells from experiencing toxic oxygen poisoning [29]. When 0.6% (vol/vol) H_2_O_2_ was added, the concentration of hydrogen peroxide in the culture medium was greater than a certain value (25 µM), and *Escherichia coli* could induce resistance to oxidative stress, with catalase becoming the main hydrogen peroxide scavenger [15, 30]. Catalase was significantly increased in *E. coli* BL21(DE3)^∆slp^ compared to wild-type *E. coli* BL21(DE3) (Fig 5), indicating that *E. coli* BL21(DE3)^∆slp^ can induce catalase to resist the harmful hydrogen peroxide environment. The mutation of the *slp* gene caused a marked (p<0.05) decrease in the glutathione S-transferase activity (Fig 6A), while it caused significantly (p<0.05) increased glutathione peroxidase activity in *Escherichia coli* (Fig 6B). An increase in catalase activity may contribute to an increase in peroxidase activity, although glutathione is not necessary for *E. coli* resistance to oxidative stress. Glutathione also may protect cells from radiation damage during oxidative stress. Thus, due to the mutation of the *slp* gene, highly expressed catalase and glutathione were synergistic in decomposing the hydrogen peroxide.

Another destruction system of H_2_O_2_ is the catalase family. In detecting catalase encoding genes, the expression of the *katE* gene did not change significantly (Fig 8). HPII is not peroxide inducible, and its gene, *katE*, is transcribed at the transition from the exponential phase to the stationary growth phase [16, 31]. At the same time, the expression level of the *ahpCF* gene encoding the alkyl hydroperoxide reductase system, which was initially characterized as rapidly reducing diverse organic hydroperoxides, was not changed (Fig 8). High extracellular levels of H_2_O_2_ diffuse into the cell, and the scavenging ability of *ahpCF* can be overwhelmed [2]. *Escherichia coli* can activate major regulatory factors in the body to perceive hydrogen peroxide, such as the OxyR regulator. HPI (*katG*), which requires the positive transcriptional activator OxyR, was transcriptionally induced during the exponential phase in response to low micromolar concentrations of H_2_O_2_ [15, 32]. The markedly high expression of the *katG* gene (Fig 8), indicating that OxyR was directly oxidized by 0.6% (vol/vol) H_2_O_2_, leads to disulfide bond formation and OxyR regulator activation of HPI to resist hydrogen peroxide oxidative stress. Cells were resistant to oxidative stress in the hydrogen peroxide environment, and the degree to which the cells were attacked by free radicals was significantly reduced (Fig 7).

## Acknowledgement

The research was financially supported by The Importation and Development of High-Caliber Talents Project of Beijing Municipal Institutions (CIT&TCD20140315).

